# Association between activities outside work and presenteeism among Korean wage workers: using nationwide survey

**DOI:** 10.1101/373241

**Authors:** SungWon Jung, June-Hee Lee, Kyung-Jae Lee

## Abstract

**Introduction:** Presenteeism is a significant global health issue nowadays and can influence to producitivity loss. So, there were many studies to analyse relationship between workplace factor and presenteeism. But a few studies have considered non-occupational factor. The purpose of this study was to examine the association between presenteeism and activities outside work, such as, volunteering, self-development, leisure/sports, gardening & house repair activity, in Korean wage workers.

**Methods:** This study analysed the 4^th^ Korean Working Condition Survey(KWCS) and overall 19,294 wage workers participated. To find the relationship between presenteeism and activities outside work, multivariate logistic regression analysis was used after adjusting for general and occupational characteristics.

**Results:** Among self-development, leisure/sports and gardening & house repair activities significantly increased the odds ratio of presenteeism (Odds ratio[OR] =1.33, 95% confidence interval[CI]: 1.23-1.43, OR = 1.40, 95% CI: 1.29-1.51 and OR = 1.09, 95% CI: 1.01-1.18, respectively).

**Conclusions:** Some of activities outside work were related to presenteeism in Korean wage workers. Although many of previous studies addressed positive effect of those activities for health, this study showed negative effect of activities oudside work for health. And we should consider Korean organizational culture for this reason and need more structural studies to find out specific factors.

## Introduction

The modern day wage workers spend half of their lives at work. Therefore, workplace health has become an important issue, not only for the individual but also for the employer. Health problems of workers are associated with the loss of productivity, which is directly linked with the interests of employers[1-3], and the indirect economic burden caused by such productivity deterioration is reported to be greater than the economic burden caused by other illnesses [4, 5]. Presenteeism is quickly gathering attention as a moot subject when discussing productivity deterioration, and various studies are under way.

Presenteeism is a contrasting concept to Absenteeism. It can be defined as a limitation to an individual’s work efficiency despite coming into work[6], and can also be defined in a number of ways including limitations to work efficiency due to coming into work with health problems [7, 8]. There are studies that suggest the productivity and work attendance of workers with health problems decrease over time[9]. There have been other studies on presenteeism in relation to its impact on health. One such study found that workers that worked three consecutive years had a higher mobility rate of cardiovascular illnesses than workers that took time off [10]. When observing workers with experience of presenteeism, they assessed their own future health to be deficient[11, 12]. According to a Swedish research, at least one third of workers experience presenteeism[8]. While in South Korea, the percentage of workers with experience of presenteeism is 21% according to the Third Korean Working Condition Survey (KWCS); presenting workplace health as an important issue[13].

Presenteeism is largely affected by organizational and personal factors[14]. A body of research has been conducted to confirm the relationships between presenteeism and organizational factors, such psychological, occupational factors in the workplace[15-17], and individual health risk factors such as smoking, alcohol drinking and physical exercise[18, 19]. Activities outside work, such as physical exercise, is a great opportunity to restore an individual’s fitness and it changes the individual health risk factor[20].

Typical examples of activities outside work are volunteering, self-development activities, gardening, house repair, cultural activities and sports[21]. Activities such as leisure sports, physical exercise and gardening may diminish symptoms of depression and anxiety, while improving quality of sleep and other psychological factors[20, 22, 23].

Only a handful of research examining the relationship between activities outside work and presenteeism has been conducted, and the existing research mainly focuses on only the relationship between physical exercise and presenteeism[24, 25]. Therefore, this research will utilize the Fourth KWCS to examine the relationship between various types of activities outside work undertaken by South Korean wage workers and presenteeism.

## Methods

### Study Subjects

This research utilized resources from the Fourth Korean Working Condition Survey, conducted by the Korea Occupational Health and Safety Agency (KOSHA) in 2014. The KWCS was developed based on the European Working Conditions Survey. The KWCS is a national open source date with safeguards to protect the participants` anonymity and privacy rights. By this reason, our study is not applicable for IRB.

Subjects of the survey were employees over the age of 15, and a total of 50,007 people were interviewed and surveyed for the Fourth KWCS. Participants who gave inadequate or incomplete answers such as ‘I don’t know/No Answer’ or refused to take part were removed from the survey results. The study is aimed towards wage workers, thus, responses from self-employed entrepreneurs, business owners, unpaid workers in family businesses and other ineligible subjects were removed. The number of people that identified themselves as soldiers, agricultural workers, forestry workers and fishery workers were very small, and was removed. Underage survey participants (15 to under 20), were also removed. Finally, participants with pre existing injuries, experience of harmful accidents, and previous cardiovascular issues that may influence an individual’s experience of presenteeism were removed. In total, the research used data from 19,294 wage earners above the age of 20.

## Measurements

### General characteristics

General characteristics included gender, age (20-29 years, 30-39 years, 40-49 years, 50-59 years, ≥60 years), level of education (Below Middle School level, High School Diploma, University Graduate and Other).

### Occupational characteristics

Occupational characteristics included employment status (regular work, temporary work or day labour), Occupation type (management/professional, office work, service/sales, technical, or simple labor), working hours per week(≤40, 41–59, ≥60h), shift work, number of employees(<50,50–299, or ≥300) and monthly income (<1,300,000 Korean won (KRW), 1,300,000–1,990,000 KRW, 2,000,000– 2,990,000 KRW, or ≥3,000,000 KRW)[26].

### Activities outside work

Items to assess ‘Activities outside work’ acted as independent variables for this research. Participants had to respond to the question “In general, how often are you involved in any of the following activities outside work?”. Activities outside work included volunteer work, self-development, sports, physical exercise, leisure activities, gardening and house repair. Answers including “more than an hour every day,” “less than an hour every day or two,” “once or twice a week,” “once or twice a month,”; or “;once or twice a year” were considered to have “taken part” in an activity outside work[26].

### Presenteeism

Presenteeism can be defined as a phenomenon of workers coming to work, even when they need to rest at home due to illness or injury[8]. Thus, the study identified participants with presenteeism, as a dependable variable, when a person answered ‘yes’ to the question "Over the past 12 months, did you work at least one day when you were sick?”.

### Data analysis

A chi-square test was conducted to determine the distribution based on the general and occupational characteristics of wage workers who have experience presenteeism. Another chi-square test was conducted to determine the distribution of presenteeism in relation to activities outside work. Next step was to analyze the correlation between activities outside work, as a independent variable, and presenteeism, a dependant variable. General and occupational characteristics were revised and run through a multiple logistic regression. Level of significance was 0.05 and all static analysis was conducted by using SPSS 14.0.

## Results

### Distribution of presenteeism according to general and occupational characteristics “Table 1”

A series of chi-square tests were used to determine the distribution of presenteeism according to general and occupational characteristics. The results found that females (23.4%) had more experience with presenteeism than males. Wage workers between the age of 40-49 (23.9%) had more experience with presenteeism than other age groups. Workers in professional and managerial jobs (23.6%) had the most experience with presenteeism among occupational types, and full-time workers (22.1%) experienced more presenteeism than temporary/non-permanent workers (18.5%). Those who worked over 60 hours a week showed more experience with presenteeism (26.3%), and the results showed that presenteeism increased according to the amount of hours worked. Percentage of workers working in a workplace of 50-299 that had experience presenteeism was 22.6%, which was comparable higher than those who worked at a workplace of under 50 staff and those who had worked at a workplace of over 300 staff, respectively. There were no statistically significant difference between level of education and shift work in relation to presenteeism.

**Table 1.**
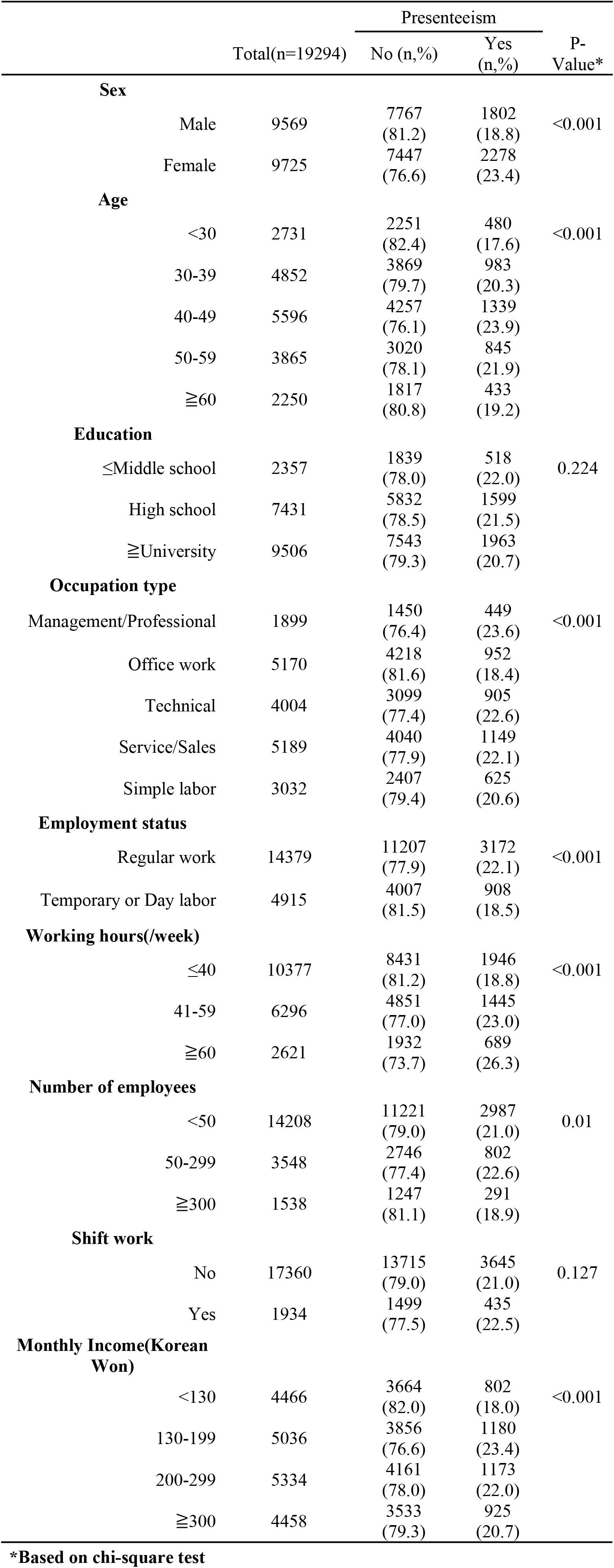
Number of workers with presenteeism by general and occupational characteristics

|  |  | Total(n=19294) | Presenteeism |  | P-Value* |
| --- | --- | --- | --- | --- | --- |
|  |  |  | No (n,%) | Yes (n,%) |  |
| **Sex** |  |  |  |  |  |
|  | Male | 9569 | 7767<br>(81.2) | 1802<br>(18.8) | <0.001 |
|  | Female | 9725 | 7447<br>(76.6) | 2278<br>(23.4) |  |
| **Age** |  |  |  |  |  |
|  | <30 | 2731 | 2251<br>(82.4) | 480<br>(17.6) | <0.001 |
|  | 30-39 | 4852 | 3869<br>(79.7) | 983<br>(20.3) |  |
|  | 40-49 | 5596 | 4257<br>(76.1) | 1339<br>(23.9) |  |
|  | 50-59 | 3865 | 3020<br>(78.1) | 845<br>(21.9) |  |
|  | ≥60 | 2250 | 1817<br>(80.8) | 433<br>(19.2) |  |
| **Education** |  |  |  |  |  |
|  | ≤Middle school | 2357 | 1839<br>(78.0) | 518<br>(22.0) | 0.224 |
|  | High school | 7431 | 5832<br>(78.5) | 1599<br>(21.5) |  |
|  | ≥University | 9506 | 7543<br>(79.3) | 1963<br>(20.7) |  |
| **Occupation type** |  |  |  |  |  |
|  | Management/Professional | 1899 | 1450<br>(76.4) | 449<br>(23.6) | <0.001 |

|  |  |  |  |  |
| --- | --- | --- | --- | --- |
| Office work | 5170 | 4218<br>(81.6) | 952<br>(18.4) |  |
| Technical | 4004 | 3099<br>(77.4) | 905<br>(22.6) |  |
| Service/Sales | 5189 | 4040<br>(77.9) | 1149<br>(22.1) |  |
| Simple labor | 3032 | 2407<br>(79.4) | 625<br>(20.6) |  |
| **Employment status** |  |  |  |  |
| Regular work | 14379 | 11207<br>(77.9) | 3172<br>(22.1) | <0.001 |
| Temporary or Day labor | 4915 | 4007<br>(81.5) | 908<br>(18.5) |  |
| **Working hours(/week)** |  |  |  |  |
| ≤40 | 10377 | 8431<br>(81.2) | 1946<br>(18.8) | <0.001 |
| 41-59 | 6296 | 4851<br>(77.0) | 1445<br>(23.0) |  |
| ≥60 | 2621 | 1932<br>(73.7) | 689<br>(26.3) |  |
| **Number of employees** |  |  |  |  |
| <50 | 14208 | 11221<br>(79.0) | 2987<br>(21.0) | 0.01 |
| 50-299 | 3548 | 2746<br>(77.4) | 802<br>(22.6) |  |
| ≥300 | 1538 | 1247<br>(81.1) | 291<br>(18.9) |  |
| **Shift work** |  |  |  |  |
| No | 17360 | 13715<br>(79.0) | 3645<br>(21.0) | 0.127 |
| Yes | 1934 | 1499<br>(77.5) | 435<br>(22.5) |  |
| **Monthly Income(Korean Won)** |  |  |  |  |
| <130 | 4466 | 3664<br>(82.0) | 802<br>(18.0) | <0.001 |
| 130-199 | 5036 | 3856<br>(76.6) | 1180<br>(23.4) |  |
| 200-299 | 5334 | 4161<br>(78.0) | 1173<br>(22.0) |  |
| ≥300 | 4458 | 3533<br>(79.3) | 925<br>(20.7) |  |

\*Based on chi-square test

### Distribution of presenteeism according to activities outside work “Table 2”

Chi square tests were conducted to determine the distribution of sleep disorders, depression and anxiety according to leisure and social activities. Experience of presenteeism was higher in those who engaged in self-development activity (23.3%) compared to those who didn’t. Presenteeism in subjects who engaged in leisure & sports activities and gardening & house repair were higher than those that didn’t (22.6% and 22.3, respectively). The results did not show any significant variances for those who engaged in volunteering activity.

**Table 2.** Distribution of presenteeism by activities outside work

|  |  | Presenteeism |  | P-Value* |
| --- | --- | --- | --- | --- |
|  | Total(n=19294) | No (n,%) | Yes (n,%) |  |
| **Volunteering** |  |  |  |  |
| No | 16837 | 13274<br>(78.8) | 3563<br>(21.2) | 0.892 |
| Yes | 2457 | 1940<br>(79.0) | 517<br>(21.0) |  |
| **Self-development** |  |  |  |  |
| No | 12543 | 10038<br>(80.0) | 2505<br>(20.0) | <0.001 |
| Yes | 6751 | 5176<br>(76.7) | 1575<br>(23.3) |  |
| **Leisure/Sports** |  |  |  |  |
| No | 6711 | 5481<br>(81.7) | 1230<br>(18.3) | <0.001 |
| Yes | 12583 | 9733<br>(77.4) | 2850<br>(22.6) |  |
| **Gardening & House repair** |  |  |  |  |
| No | 13308 | 10564<br>(79.4) | 2744<br>(20.6) | <0.001 |
| Yes | 5986 | 4650<br>(77.7) | 1336<br>(22.3) |  |

\*Based on chi-square test

### Relationship between activities outside work and presenteeism “Table 3”

Multivariate logistic regression analysis was performed to determine the relationship between presenteeism according to activities outside work. The previously mentioned general and occupational characteristics were adjusted and analyzed. The results showed that the risk of presenteeism was higher in those who engaged in self-development activities to those that did not (OR = 1.33 [95% CI: 1.23-1.43]). The risk was also apparent for those who engaged in Leisure & Sports activity, compared to those that didn’t (OR = 1.40 [95% CI: 1.29–1.51]). Lastly, the higher risk was also apparent in those who engaged in gardening & house repair activity, compared to those that did not (OR = 1.09 [95% CI: 1.01–1.18]).

**Table 3.** The odds ratios and 95% confidence intervals of activities outside work on presenteeism

|  | Model I* |  | Model II† |  |
| --- | --- | --- | --- | --- |
|  | OR | 95% CI | OR | 95% CI |
| **Volunteering**<br>(Ref: No) | 0.99 | 0.90 - 1.10 | 0.99 | 0.90 - 1.11 |
| **Self-development**<br>(Ref: No) | 1.22 | 1.14 - 1.31 | 1.33 | 1.23 - 1.43 |
| **Leisure/Sports**<br>(Ref: No) | 1.31 | 1.21 - 1.41 | 1.40 | 1.29 - 1.51 |
| **Gardening & House**<br>**repair**<br>(Ref: No) | 1.11 | 1.03 - 1.19 | 1.09 | 1.01 - 1.18 |

Calculated using multiple logistic regression analysis
\*Crude odds ratio
†Adjusted for age, sex, education, employment status, number of employees, working hours, shift work, monthly income

## Discussions

This study was able to examine relationships between the average South Korean wage workers’ activities (volunteering, self-development, leisure & sports activities, gardening & house repairs) and the degree of presenteeism. The study found that the risk of presenteeism was notably high in individuals that engaged in self-development, leisure & sport activities or gardening & house repairs. Most research utilizing activities outside work as a variable reported positive results, and many studies examined the effects of sports, leisure time and physical activity to health. It is well established that engaging in leisure time, physical activity and sports can lower the risk of obesity[27] and cardiovascular diseases[28]. Past studies have shown how obesity and cardiovascular diseases can raise the risk of presenteeism[29]. We expected leisure and sports activities to lower the risk of presenteeism, as indicated by previous research, but the study resulted in the opposite.

In previous research, leisure time and physical activity outside of work coined with high levels of occupational physical activity was shown to have harmful effects on mortality and risk of cardiovascular diseases[30, 31]. South Korean workers are among one of the most worked in the OECD, and are known for their high rates of occupational stress and work intensity. In a previous study on South Korean wage workers, over 70% of workers were exposed to occupational physical activity such as painful or tiring positions, repetitive hand or arm movements, moving or lifting heavy loads and standing posture, and these workers were more susceptible to presenteeism[13]. We believe these factors. There is a possibility that these factors may have driven our research results. Also, involuntary leisure time and physical activity embedded into South Korea’s work culture such as company sports days, weekend hikes and other after work activities may have also contributed to the results.

This study showed that engaging in self-development activities increased the risk of presenteeism. A previous study that used the Fourth KWCS found that self-development activities increased the risk of sleep disorders[26]. The study concluded that self-development activities undertaken by South Korean workers were not for self-satisfaction but rather a tool to gain promotion or secure employment, which can be a cause of stress. Another reason was that because such activities occur outside work hours, it reduces time for rest[32]. There are also studies that identified a group suffering from insomnia and sleep disorders to be vulnerable to presenteeism[33], and these results correlate with our own research findings. The implications of such finding may suggest that South Korean wage workers aren’t voluntarily engaging in leisure, sports and self-development activities, but involuntarily at the behest of others, trapping South Korean workers under the shadow of the workplace even after business hours.

This study also found that gardening & house repair activities increased the risk of presenteeism. Most previous research found gardening activity to have a positive impact on health, especially in reducing depression and anxiety and improving mental health[22]. We expected a similar result for presenteeism, but the study showed contrary findings. As a result, a close examination of South Korea’s housing culture may be needed. Most South Korean live in apartment buildings, not residential houses, and apartments are viewed as a prized asset and used to show other one’s social status and/or wealth[34]. Therefore, house repairs can be a considerable economic burden and may have negative effects such as leading to stress.

Our study has some strengths. The first strength of this study is that is the first to investigate the relationship between various activities outside work and presenteeism, and providing an opportunity to think about the health effects of activities outside work.

Secondly, after work hours of South Korean wage workers may be an extension of work, rather than the end, and the study showed the possibility of adverse health effects as result. Such findings highlight the need for additional research to closely examine domestic work environments. In addition, the European Working Conditions Survey used for the survey may need to be reconstructed into a new survey to accurately reflect the cultural specificity of the country as items can lead to different interpretations based on an individual’s cultural viewpoint.

The study has its limitations. First, as a cross-sectional study, it did not reveal the causality between activities outside work with presenteeism. Though it did present a relationship that did not previously exist, and so it was meaningful in the sense that new alternatives were presented. Second, the exclusion of personal factors such as alcohol intake and smoking as survey items meant that those items were not included in the revised general characteristic category. However, there are many existing studies based on the KWCS and the European Working Conditions Survey, and so the value of this study is still sufficient.

Third, the definition of presenteeism, which is a dependent variable, is from a self-reporting questionnaire, which may lowers it’s objectivity. However, many of the existing studies have used self -reporting questionnaires to show sufficient validity.

Despite the outlined limitations above, the research results will be a valuable resource for further research to find new solutions and alternatives to the distinct work environment of South Korea’s wage workers.

## Conclusions

This study was able to show that various activities outside work can increase the risk of presenteeism. The results of this study indirectly suggests that after work hours for South Korean wage workers are merely an extension of work, a unique characteristic of South Korea’s work culture, and may be a cause of presenteeism. On another note, the survey items of the European Working Conditions Survey should be translated to better reflect the cultural viewpoints of various countries. In conclusion, further research examining specific components of South Korea’s unique work culture that led to our results is necessary.

## Acknowledgements

None.

## Declarations

Ethics approval and consent to participate Not applicable

## Consent to publish

Not applicable

## Availability of data and materials

Data sharing not applicable to this article as no datasets were generated or analysed during the current study.

## Competing interest

The authors have no competing interests

## Funding

This work was supported by the Soonchunhyang University Research Fund.

## Author’s contributions

Study conception and design: SW Jung, KJ Lee, JH Lee; Data acquisition: SW Jung; Data analysis and interpretation: KJ Lee, SW Jung, JH Lee; Drafting the manuscript: SW Jung; Critical revision: KJ Lee, JH Lee. All authors read and approved the final manuscript.

